# DLPacker: Deep Learning for Prediction of Amino Acid Side Chain Conformations in Proteins

**DOI:** 10.1101/2021.05.23.445347

**Authors:** Mikita Misiura, Raghav Shroff, Ross Thyer, Anatoly B. Kolomeisky

**Affiliations:** Department of Chemistry, Center for Theoretical, Biological Physics, Rice University, Houston, TX 77005; CCDC Army Research Lab—South, Austin, TX 78712; Department of Chemical and Biomolecular Engineering, Rice University, Houston, TX 77005; Department of Chemistry, Department of Physics and Astronomy, Department of Chemical and Biomolecular Engineering, Center for Theoretical Biological Physics, Rice University, Houston, TX 77005

**Keywords:** 3DCNN, DNN, U-net, Protein Structure Prediction, Side Chain Restoration

## Abstract

Prediction of side chain conformations of amino acids in proteins (also termed ‘packing’) is an important and challenging part of protein structure prediction with many interesting applications in protein design. A variety of methods for packing have been developed but more accurate ones are still needed. Machine learning (ML) methods have recently become a powerful tool for solving various problems in diverse areas of science, including structural biology. In this work we evaluate the potential of Deep Neural Networks (DNNs) for prediction of amino acid side chain conformations. We formulate the problem as image-to-image transformation and train a U-net style DNN to solve the problem. We show that our method outperforms other physics-based methods by a significant margin: reconstruction RMSDs for most amino acids are about 20% smaller compared to SCWRL4 and Rosetta Packer with RMSDs for bulky hydrophobic amino acids Phe, Tyr and Trp being up to 50% smaller.

## 1 Introduction

*De novo* protein structure prediction is one of the key fundamental problems in structural biology and a lot of research efforts are currently focused in this area. Accurate prediction of the protein structure from its amino acid sequence is required in order to understand the molecular details of various biological phenomena and to fully uncover biological functions of large number of proteins found inside genomes of living organisms. It can enable efficient *in silico* protein design, allowing for the development of new enzymes catalyzing novel types of reactions, development of new target-specific protein therapeutics and many other powerful applications. However, despite the fact that a protein’s fold, in principle, is fully defined by its amino acid sequence, the prediction task is extremely complex due to the immense size of the conformational search space, which grows exponentially with the sequence length.

Existing approaches often split protein structure prediction into two steps: prediction of the protein’s backbone conformation (‘folding’) and prediction of the side chain conformations (‘packing’). Most of the currently available methods for side chain packing are physics-based approaches that involve some sort of search inside a given sample space, often defined by a library of pre-defined rotamers and/or an optimization using a well-designed energy function. [1, 2, 3, 4, 5, 6] These methods view the problem from a physico-chemical perspective and are trying to optimize interactions between side chains while avoiding steric clashes and minimizing the overall energy of the system.

There are currently multiple breakthroughs in Machine Learning (ML) methods that are revolutionizing multiple areas of science, including chemistry, physics, biology and medicine. A large number of recent studies explored the potential of applying Neural Networks (NNs) and Deep Neural Networks (DNNs) for a variety of difficult problems for which classical methods are still unsuccessful. This includes protein modeling, small molecule property predictions, drug discovery and materials design. [7, 8, 9, 10, 11, 12, 13, 14, 15, 16, 17] One of the most notable works in the field of protein structure prediction is AlphaFold and its successor AlphaFold2, which significantly outperform classical methods for protein structure prediction. [8, 18]

ML methods have already been used for the task of amino acid side chain restoration. [12, 19, 20, 21, 22] Nagata et al. trained an ensemble of 156 shallow NNs using inter-atomic distances as an input, which resulted in a remarkably fast side chain restoration algorithm. The latest version of the OPUS side-chain modeling framework, named OPUS-RotaNN, which is based on OPUS-TASS, uses deep neural networks to achieve even better performance. [19] OPUS-RotaNN significantly improves rotamer sampling and can outperform other methods like OSCAR-star, SCWRL4 and FASPR.

In this work we explore the potential of DNNs for the protein side chain conformation prediction. The problem is formulated as an image-to-image transformation and a U-net-style DNN is utilized to solve it. We demonstrate that our method significantly outperforms other published approaches. Reconstruction RMSDs for most amino acids are about 20% smaller compared to SCWRL4 and Rosetta Packer, and RMSDs for bulky hydrophobic amino acids Phe, Tyr and Trp are up to 50% smaller. We anticipate that our approach can be useful for a variety of structural biology-related tasks such as homology modeling, generating point mutants for *in silico* protein engineering and other related problems. Since the direct-coupling analysis (DCA) methods are successful in predicting contacts between amino acids and inferring the backbone conformations of proteins, it may be possible to couple our method with DCA to deliver high quality full atom structure predictions. [23] Our code is available at: https://github.com/nekitmm/DLPacker

## 2 Methods

When training a 3DConv Neural Network model to solve the problem of side chain packing there are three major steps: input generation, NN inference and side chain reconstruction. The overall approach is schematically illustrated in Figure 1. The next three subsections describe these steps in more detail.

**Figure 1:**
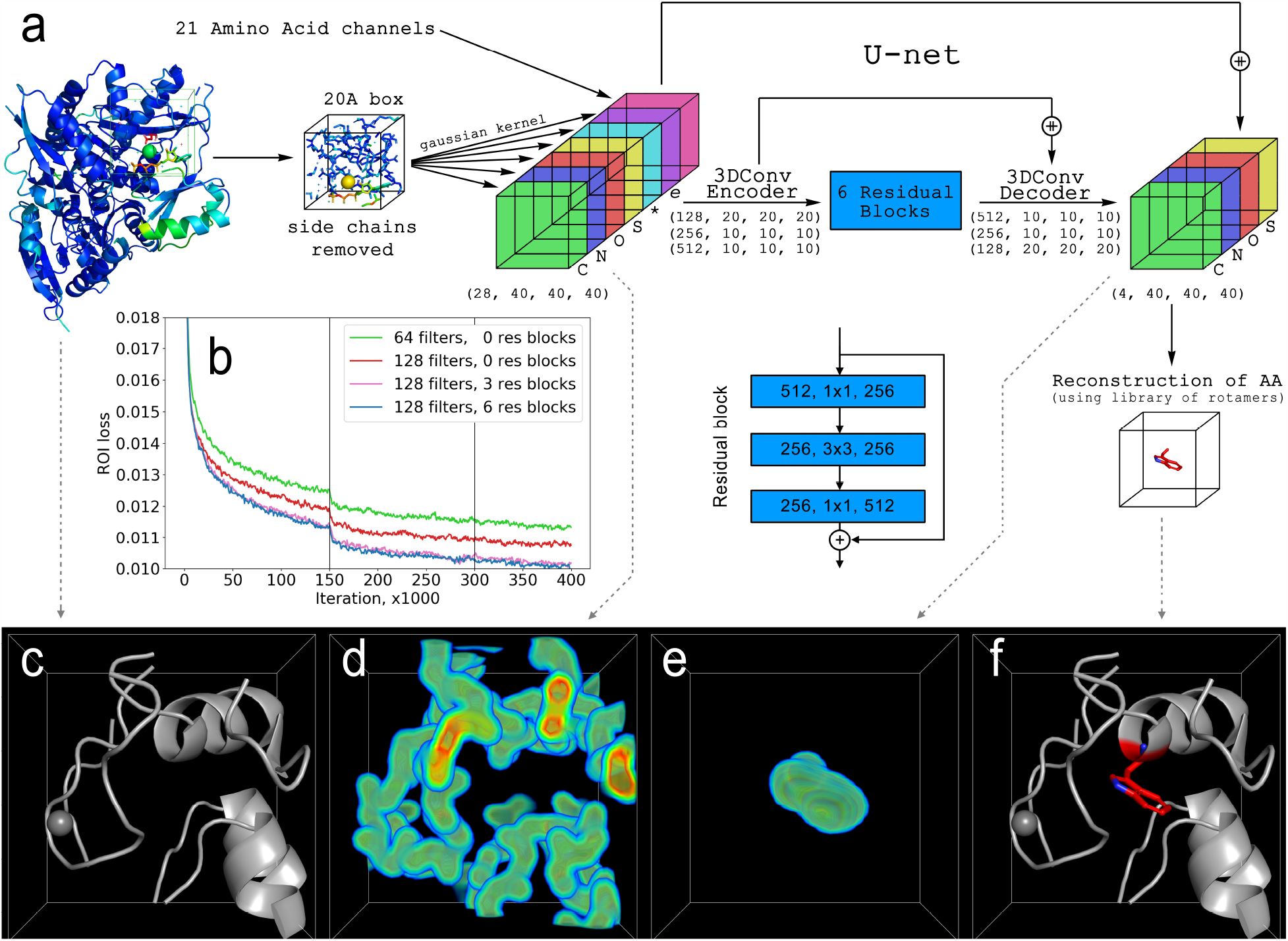
Overview of our side chain restoration algorithm. **a**. Input data generation and U-net architecture. After an amino acid is chosen, a cube comprising it’s microenvironment is isolated from the protein and atom positions are placed on a grid and split into a number of different input channels. The input is then fed into the 3DConv U-net, consisting of encoder, six residual blocks and a decoder. Structure of a residual block is shown in blue. The output is a 3D density, outlining predicted side chain conformation. This prediction is then turned into side chain atom positions by fitting a pre-calculated library of side chain conformations. Vertical lines correspond to step-wise decreases of the learning rate. **b**. Loss curves for different NN architectures with different numbers of filters in the first 3DConv layer and different numbers of residual blocks. **Bottom row:** Illustration of input and output. **c**. Cartoon representation of the amino acid’s microenvironment. Only the backbone is left at this stage. **d**. Encoded atom positions. Each atom is represented as a 3D gaussian density kernel. Only the sum of four element channels (C, N, O, S) is shown for simplicity. **e**. Output from the U-net. Sum of four element channels (C, N, O, S) is shown. **f**. Reconstructed amino acid side chain (Trp in this case, shown in red).

### 2.1 Description of the Dataset Preparation

Training and validation datasets were created as described by Shroff et al. [24] In short, all entries in the PDB database were clustered to 50% similarity to avoid bias towards more abundant classes of proteins. From each cluster only the single structure with the highest resolution was selected and remodeled using the PDB-redo algorithm, which rebuilds all structures using a unified algorithm. [25] Clusters containing only structures with resolution lower than 2.5Å were discarded. If multiple alternative conformations exist inside the protein structure, only the alternative conformation with the largest sum of occupancies was used, and other conformations are discarded. Hydrogen atoms were omitted in our model and selenomethionine residues were converted into methionine.

This process yielded 19,436 structures, of which 2,201 were reserved as a validation set and the rest were used as a training set. This resulted in about 10 million training examples and one million validation examples.

### 2.2 Input Data Generation

The process of input generation is schematically described on the top portion of Figure 1. After a target amino acid is chosen for side chain reconstruction, a 20 Å box containing its surrounding microenvironment is isolated from the protein. The side chain of the target amino acid (if present) is always removed from the input. For all other side chain atoms in the box, there are different options depending on what stage (training/validation/inference) is being performed by the model. Since the neural network in this setup is used to restore all side chains in a sequential manner (one amino acid at a time), the method should be able to make predictions using both a pure backbone representation of a protein and a partially restored representation of a protein (with some of the side chains already present). Therefore, during the training stage the side chains of non-target amino acids were removed using the following algorithm: in 50% of training examples all side chains were removed from the input and in the other 50% a random fraction of the side chains were removed. All of the amino acid side chains in the PDB structure were completely removed prior to the validation stage and then restored one by one.

After the input box is defined and side chains are removed if necessary, all atoms are placed on a grid with 40×40×40 dimensions and split into 28 channels: five element channels (one channel each for **C, N, O, S** and one for all other elements), one partial charge channel (partial charges were taken from the Amber99sb force field), 21 amino acid channels (one for each of the 20 canonical amino acids plus one for all other amino acids), and one label channel that encodes an amino acid label to restore. While the purpose of all element channels and the partial charge channel is clear, the purpose of 21 amino acid channels is to propagate the information about the sequence, which is necessary because some or even all side chains are removed. The *backbone atoms* of each amino acid in the input box are *repeated* in the respective channel (in addition to being present in their respective element channels). For example, if there is a phenylalanine residue present, its backbone (**C, N, CA** and **O**) atoms are copied into one phenylalanine channel. This way the information about protein sequence is not lost after removal of side chains. The output is similar to the input with two major differences: only four element channels are present (**C, N, O** and **S** since only these four elements are present in amino acid side chains) and the *target* side chain is present. Note that only one *target* side chain is restored at a time, while all the other side chains present in the box are left as they are.

### 2.3 Neural Network Architecture and Training

We conceptualized the problem of side chain restoration as an image-to-image transformation and opted for a U-net architecture which is frequently used in this field. [26, 27, 28, 29, 30] One major difference, however, is that our inputs are 3D images, which require the use 3D convolutions in our architecture. The final architecture is shown on the right in Figure 1. First, amino acid labels are added as an additional channel through a fully connected (FC) layer, whose output is then reshaped and concatenated with the rest of the input. The encoder has three strided 3D convolutional layers with 3×3 kernel size, stride 2 and ReLU (Rectified Linear Unit) nonlinearities. The number of convolutional filters are 128, 256 and 512, respectively. After the encoder, six residual blocks follow with identical architecture shown in the middle of Figure 1. Residual blocks also use ReLU nonlinearities between convolutions. The final part is the decoder, which essentially mirrors the structure of the encoder, but uses 2x upsampling to scale tensor sizes up and Leaky ReLUs as nonlinearities (*α* = 0.2). As the U-net architecture implies, the respective outputs from encoder layers were concatenated with outputs of decoder layers before feeding into the next layer. The very last 3D convolution used four filters to output the four element channels. This architecture was implemented in Python using TensorFlow 2.4. The NN was trained for about 1.5 epochs with a step-wise learning rate schedule and a batch size of 32. The initial learning rate was 1 · 10^−5^ and was decreased by a factor of 10 each time loss stopped improving. The loss was Mean Absolute Error (MAE) between prediction and ground truth. Additionally, we added what we call a Region of Interest (ROI) loss, which is just MAE in the voxels with side chain atoms being restored, which was given larger weight (100 in the final version) to force the NN to put more attention into that region.

### 2.4 Side Chain Restoration

The output from our model is not atom positions, but a 3D density map, where values of each voxel (the 3D analog of a pixel) are proportional to the probability of finding an amino acid’s side chain atom at that location (Figure 1c, bottom row). Converting this output into actual atom positions can be done in a number of ways. We settled on an approach that uses a library of side chain conformations because this approach guarantees valid conformation as a result. We used our training dataset to build a library of side chain conformations for each amino acid. The library is backbone-independent and created in such a way that it contains a list of side chain conformations in which RMSD between any individual conformation and all other conformations exceeds some threshold value. The threshold values for each amino acid were manually tuned so that the final library includes a decent, but not overwhelming number of different conformations. Threshold values that we used as well as the final numbers of side chain conformations are presented in Table 1.

**Table 1:** The side chain library used in this work. Threshold value represents minimum RMSD distance between any pair of rotamers in the library. The library was constructed from the training set. Further details are in the text.

| AA Name | Threshold, Å | Number of conformations | AA Name | Threshold, Å | Number of conformations |
| --- | --- | --- | --- | --- | --- |
| Arg | 0.50 | 2017 | Asn | 0.05 | 1228 |
| Lys | 0.25 | 2209 | Asp | 0.05 | 1066 |
| Phe | 0.05 | 1106 | Ser | 0.01 | 641 |
| Tyr | 0.05 | 1346 | Leu | 0.05 | 1329 |
| Trp | 0.05 | 4238 | Thr | 0.05 | 226 |
| His | 0.05 | 1240 | Ile | 0.05 | 677 |
| Glu | 0.25 | 903 | Cys | 0.01 | 388 |
| Gln | 0.25 | 932 | Val | 0.02 | 382 |
| Met | 0.10 | 2039 | Pro | 0.01 | 243 |

After NN inference, all side chain conformations in the library for a target amino acid are placed on a grid (see **Input Data Generation** above) and compared to the NN’s prediction by calculating a fitness score as the mean absolute difference between the prediction and the current conformation. The conformation with the lowest score is then chosen as the final prediction. An advantage of this procedure is that it guarantees the prediction to be a valid conformation. The final fitness score can also be used as a measure of prediction quality: higher scores correspond to poorer prediction quality and therefore higher uncertainty.

### 2.5 Detection of Salt Bridges

We consider that a salt bridge between Arg or Lys and Asp or Glu is formed if any of the side chain nitrogen atoms of Arg or Lys are within 3.2 Å of any side chain oxygen atoms of Asp or Glu.

### 2.6 SCWRL4 and Rosetta Packer Validation

SCWRL4 (ver 4.02) was run in default configuration. For Rosetta Packer (ver 3.12) the maximum number of rotamers was considered by passing -EX1, -EX2, -EX3 and -EX4 flags and invoking *fixbb* using the default scoring function (*ref2015*) unless stated otherwise. [31, 32] We also tried additionally increasing the number of rotamers for surface residues by using *extrachi_cutoff 0*, but that only resulted in marginal improvement of RMSDs while leading to much longer calculation times.

## 3 Results

### 3.1 Architecture of Neural Network

We first started by optimizing some of the hyperparameters of our model, including the size of the microenvironment in relation to the network input and output, number of convolutional filters in the NN and number of residual layers, presented in Figure 1b. We found that increasing the number of convolutional filters gives significant improvements in performance up to 128 filters in the first convolutional layer (the subsequent layers of decoder always double the number of filters of the previous layer), but further increases did not yield further improvements, perhaps due to larger memory consumption and smaller batch size during training. We then found that the addition of residual blocks between the encoder and decoder also leads to significant improvement in performance. We tried adding up to nine residual blocks in increments of three. Six residual blocks gave the best performance boost and further increase did not seem to improve the loss further in our experiments. Finally, we hypothesized that using a larger microenvironment (compared to our initial 20Åbox) could yield further improvements. We then tried using 30Åbox as the input and output size, but that did not lead to any measurable gains in performance, while requiring about three times longer training and predictions times. Additionally, the use of BatchNorm layers also did not give any performance boost in our experiments. Our final architecture is therefore illustrated in Figure 1.

### 3.2 Performance Analysis

Figure 1 illustrates our best performing architecture. We found that residual blocks in the middle significantly boost performance, with six blocks delivering the best performance. Further increase in the number of blocks did not lead to further improvement in our experiments. Increasing the number of convolutional filters also improves the performance at the cost of longer training and inference times. BatchNorm layers also did not give any performance boost in our experiments.

We used RMSD as a key metric to measure the performance of our algorithm and to compare it to existing models for amino acid side chain prediction. Our method’s performance is compared with that of SCWRL4 and Rosetta Packer, as both are well known and widely utilized approaches. [1, 33, 34, 35, 36, 31, 37] Each algorithm was used on ∼1,000 PDB structures from our validation dataset. The results of these comparisons are presented in Figure 2 and Table 2.

**Figure 2:**
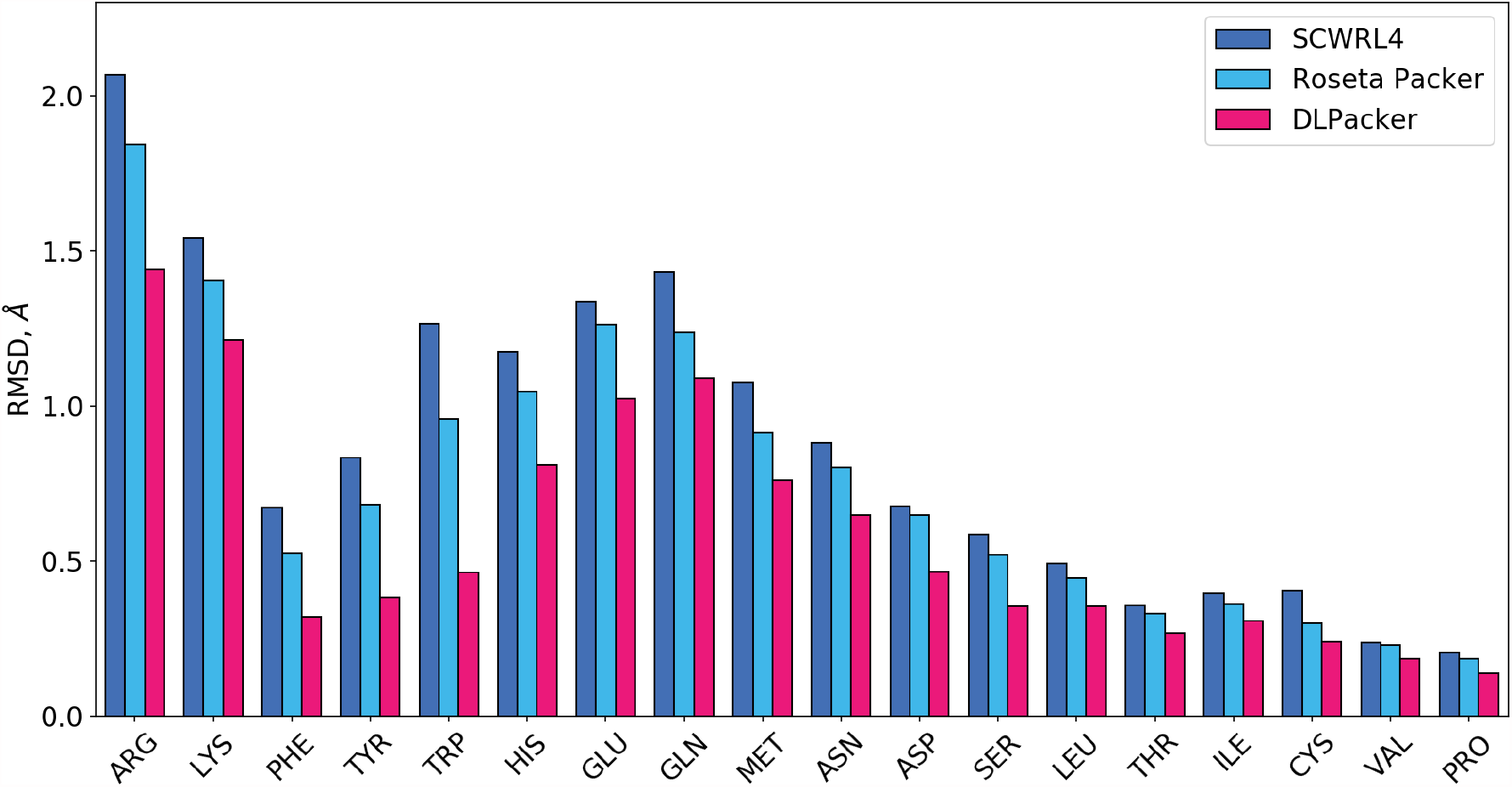
Side chain prediction RMSDs for SCWRL4, Rosetta Packer and our DLPacker (lower is better). The values for each amino acid are obtained by averaging RMSD values for ∼1,000 PDB structures from the validation set. DLPacker outperforms other methods on every amino acid.

**Table 2:** Comparison between validation RMSD values for SCWRL4, Rosetta Packer and DLPacker. The values for each amino acid are obtained by averaging RMSD values for ∼1,000 PDB structures from the validation set.

| AA Name | Number of residues | RMSD, Å |  |  |
| --- | --- | --- | --- | --- |
|  |  | SCWRL4 | Rosetta Packer | DLPacker |
| Arg | 22200 | 2.07 | 1.84 | 1.44 |
| Lys | 23781 | 1.54 | 1.40 | 1.21 |
| Phe | 17450 | 0.67 | 0.53 | 0.32 |
| Tyr | 14778 | 0.83 | 0.68 | 0.38 |
| Trp | 5754 | 1.27 | 0.96 | 0.46 |
| His | 9714 | 1.18 | 1.05 | 0.81 |
| Glu | 30014 | 1.34 | 1.26 | 1.02 |
| Gln | 15445 | 1.43 | 1.24 | 1.09 |
| Met | 9663 | 1.08 | 0.91 | 0.76 |
| Asn | 17319 | 0.88 | 0.80 | 0.65 |
| Asp | 24531 | 0.68 | 0.65 | 0.47 |
| Ser | 25371 | 0.59 | 0.52 | 0.36 |
| Leu | 41268 | 0.49 | 0.45 | 0.35 |
| Thr | 22931 | 0.36 | 0.33 | 0.27 |
| Ile | 24748 | 0.40 | 0.36 | 0.31 |
| Cys | 5377 | 0.40 | 0.30 | 0.24 |
| Val | 31443 | 0.24 | 0.23 | 0.19 |
| Pro | 19055 | 0.21 | 0.19 | 0.14 |

As one can see from Figure 2, DLPacker significantly outperforms both SCWRL4 and Rosetta Packer for all amino acids. The most significant improvement is achieved for large hydrophobic amino acids like Trp, Phe and Tyr. The improvement is also significant for a number of charged and polar amino acids such as Arg, Lys, His, Asp and Glu.

To evaluate the performance of DLPacker in more detail, we first analysed RMSD histograms for bulky hydrophobic amino acids Phe, Tyr, Trp and His (Figure 3). Histograms for all four amino acids show two major peaks: one peak in the region of low RMSDs (below 2 Å) and one in the region of high RMSDs (2 Å and above). The first peak represents correct prediction of *χ*_1_ dihedral angle, which leads to small RMSD errors during reconstruction. Small RMSD errors equate to correct reconstruction of the side chain and that the entire side chain will be located in approximately the correct location. Here we say ‘approximately in the correct location’ because it is important to take into account the existence of thermal fluctuations at normal temperatures, the limited resolution of experimentally determined structures, and other factors that lead to impossibility of precise reconstruction. The second peak, however, represents incorrect prediction of *χ*_1_, which places side chains in the wrong place entirely. This scenario is problematic as it not only leads to an incorrect reconstruction of one amino acid, but can potentially lead to incorrect reconstruction of the entire surrounding microenvironment.[38, 39] Based on this observation, we measured what we call error rates for all three algorithms and compare them in Table 3. It can be seen that DLPacker significantly reduces error rates for all four bulky hydrophobic amino acids. For example, in the case of Phe, the error rate decreases from 4.8% for SCWRL4 and 3.6% for Rosetta Packer to 1.2% for DLPacker. Similar improvements are observed for Tyr and Trp: DLPacker demonstrates about a 4-fold reduction in error rate compared to SCWRL4 and a 3-fold reduction compared to Rosetta Packer. The improvement for His is more moderate: about 1.8-2 times.

**Table 3:** Side chain reconstruction error rates for bulky amino acids. Values are measured on ∼1,000 PDB structures from the validation set. Errors are defined as predictions with high RMSDs, higher than the particular threshold. The motivation for this definition is illustrated in Figure 3 and addressed in more detail in the text. DLPacker demonstrated smaller error rates resulting in higher quality of reconstruction of side chains of bulky amino acids.

| AA Name | Threshold, Å | Error Rate, % |  |  |
| --- | --- | --- | --- | --- |
|  |  | SCWRL4 | Rosetta Packer | DLPacker |
| Phe | 3.0 | 4.8 | 3.7 | 1.2 |
| Tyr | 3.0 | 6.2 | 5.0 | 1.4 |
| Trp | 2.0 | 20.3 | 14.6 | 5.1 |
| His | 2.5 | 11.1 | 10.8 | 6.1 |

**Figure 3:**
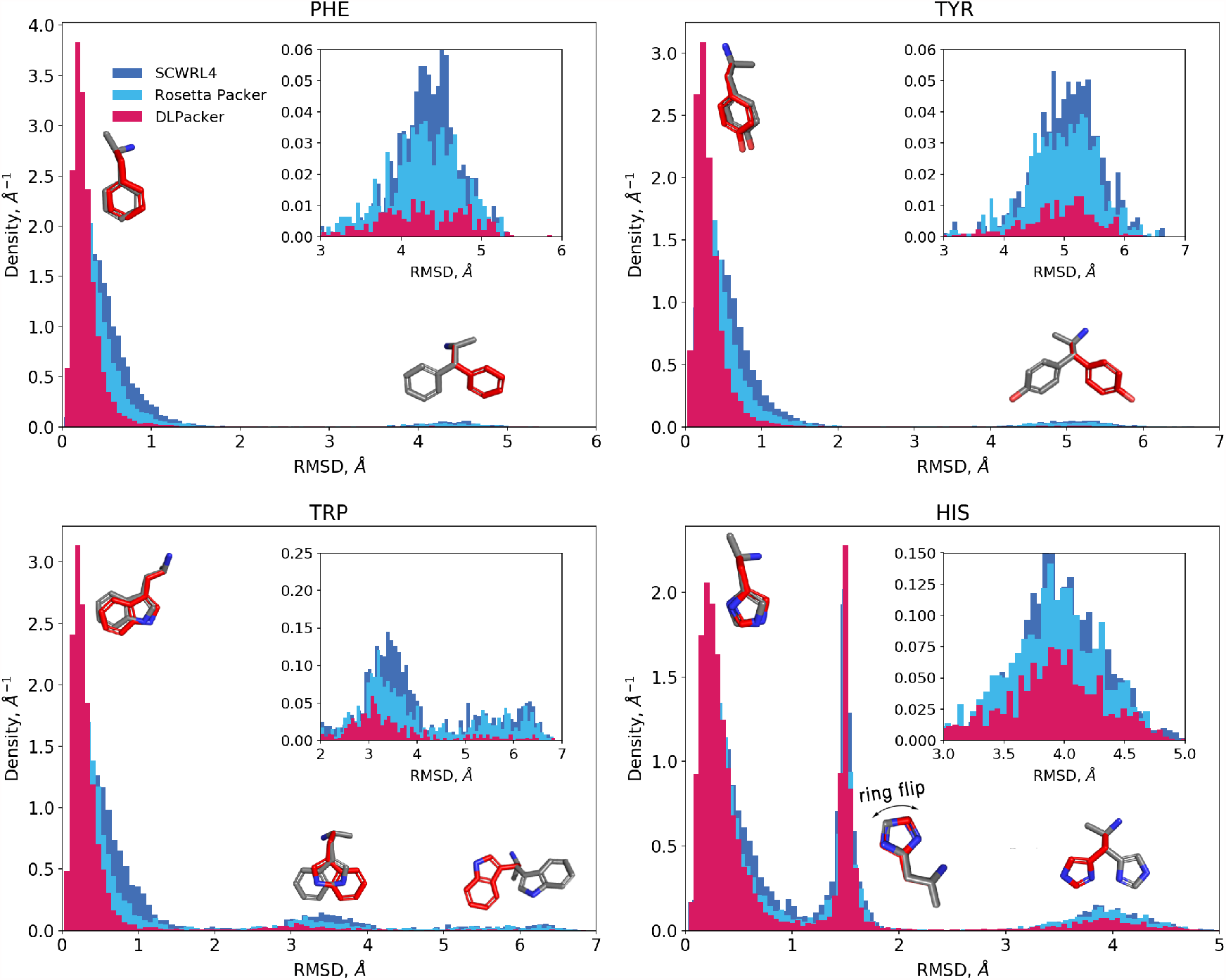
Side chain prediction RMSDs for four bulky amino acids: Phe, Tyr, Trp and His. RMSD values are measured on ∼1,000 PDB structures from the validation set. All histograms have similar structure and show two major peaks (three in the case of His, however two of the peaks can be viewed together for our purposes). Insets show high RMSD peaks in more detail. Cartoon models of side chains (red is ground truth, red is prediction) show specific examples of side chain restoration errors that compose each peak. Our algorithm (DLPacker) yields significantly fewer predictions with high RMSD resulting in higher quality packing.

Figure 4 shows how the error rates depend on an amino acid’s position in a protein. We use atom counts in an amino acid’s microenvironment as an indicator of its position within a protein: higher atom counts correspond to the denser environment of a protein’s core, while lower atom counts indicate either a very small protein or the surface of a protein. This is important since erroneous side chain predictions for bulky amino acids in dense environments are much more likely to cause incorrect packing of larger regions around them. As a result, large portions of protein’s hydrophobic core have a chance of not being reconstructed accurately, which can be problematic for some downstream tasks. In contrast, an incorrect side chain conformation on the surface of a protein might not lead to significant disruptions. As Figure 4 illustrates, the error rate for all algorithms decreases significantly as the number of atoms in the microenvironment increases, which is expected as more crowded environment leads to more interactions with neighbouring residues. DLPacker exhibits significantly smaller error rates across the whole range of atom counts and for all amino acids presented in Figure 4.

**Figure 4:**
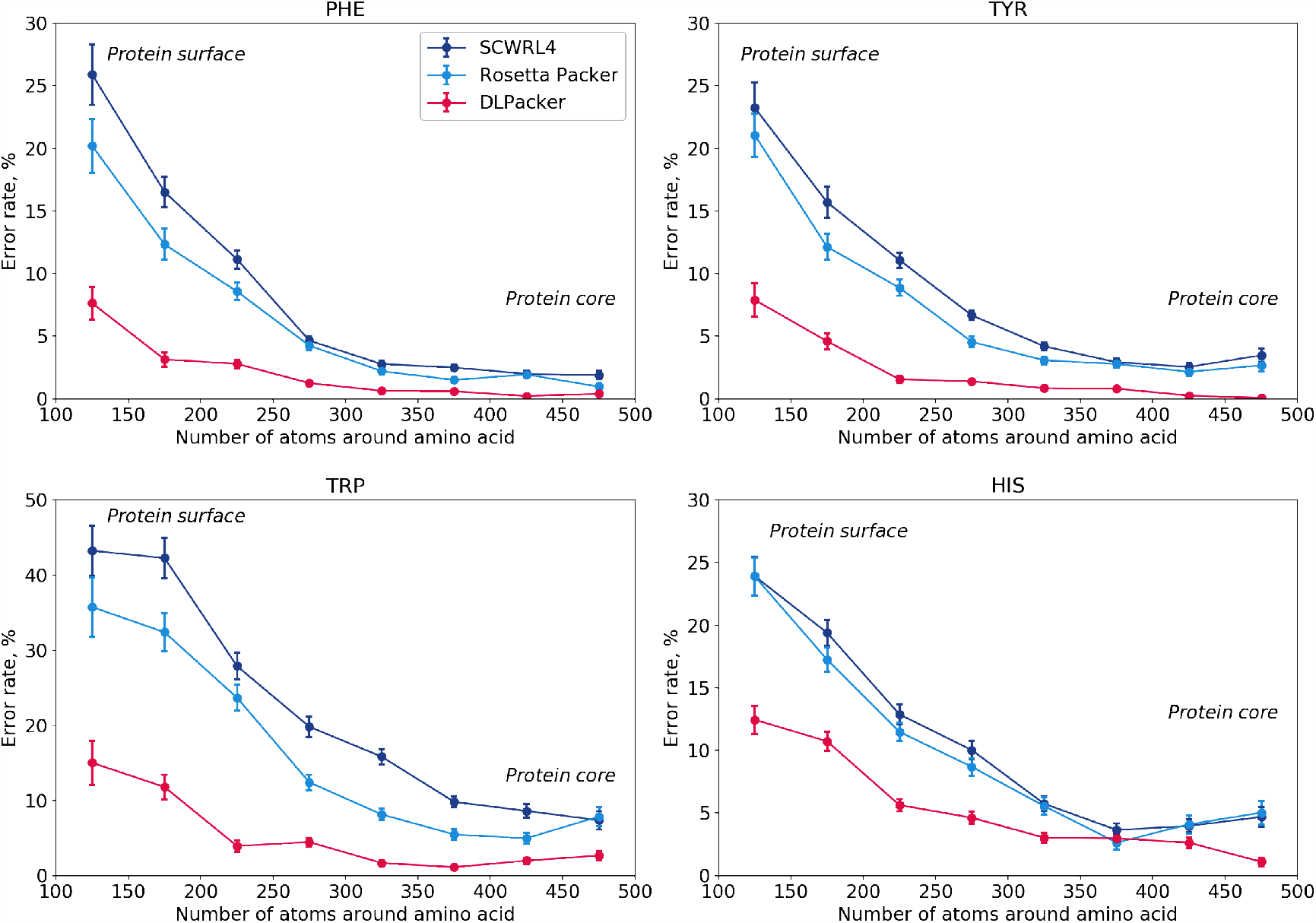
Side chain reconstruction error rates for bulky amino acids as a function of the number of atoms in their microenvironment. Values are measured on ∼1,000 PDB structures from the validation set. Vertical lines show standard deviations. The definition of the error rate is defined in detail in the text. The number of atoms in the surrounding microenvironment is an indicator of an amino acid’s position in a protein: higher atom counts indicate that an amino acid is located in the protein’s core, while low atom counts indicate a location on or near the surface. DLPacker demonstrates smaller error rates at all atom counts resulting in higher quality of reconstruction of cores of proteins as well as their surfaces.

Next, we wanted to investigate the quality of side chain restoration for charged amino acids, like Arg, Lys, Asp and Glu. Since these amino acids have multiple degrees of freedom (up to four dihedral *χ* angles), an analysis similar to that performed for bulky hydrophobic amino acids above is not possible, as the RMSD histograms do not show any useful structure. Instead, we decided to analyze the quality of reconstruction of salt bridges formed by charged amino acids since salt bridges play a significant role in protein structure formation and stabilization. Results of this analysis are presented in Table 4. We used a precision-recall metric to quantify the quality of salt bridge reconstruction. The precision value shows what fraction of salt bridges in reconstructed protein structure exist in the original PDB structure, while the recall value shows what fraction of salt bridges from the original PDB structures were reconstructed. The *F*_1_ score is a harmonic mean of precision and recall:

**Table 4:**
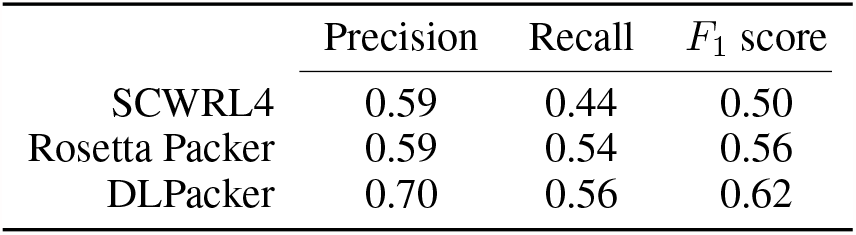
Salt bridge reconstruction metrics measured on ∼1,000 PDB structures from the validation set. DLPacker outperforms SCWRL4 and Rosetta Packer by all three metrics indicating higher quality side chain reconstruction for charged amino acids. Details are in the text.

| | Precision | Recall | $F_1$ score |
| --- | --- | --- | --- |
| SCWRL4 | 0.59 | 0.44 | 0.50 |
| Rosetta Packer | 0.59 | 0.54 | 0.56 |
| DLPacker | 0.70 | 0.56 | 0.62 |

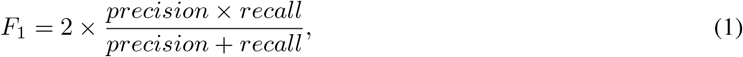

which serves as an aggregated value showing the quality of prediction and combines precision and recall into a single value. We can see that DLPacker outperforms both SCWRL4 and Rosetta Packer leading to the highest *F*_1_ score among the three: 0.62 vs 0.56 for Rosetta Packer and 0.50 for SCWRL4.

It is also interesting to look at the comparison between Mean Absolute Errors (MAE) of *χ* angles produced by the algorithms in comparison (Table 5). DLPacker works much better than other methods when comparing MAEs of *χ*_1_, but as we move to *χ*_2_, *χ*_3_ and *χ*_4_ it starts to lose its advantage. Charged and polar amino acids with lots of degrees of freedom (Lys, Arg, Glu, Gln) seem to be the most challenging ones for our method: DLPacker is often worse at predicting *χ*_3_ and *χ*_4_ than SCWRL4 and/or Rosetta Packer. Apparently, the model struggles assigning conformations to amino acids with large number of degrees of freedom (the number of available conformations grows exponentially with the number of degrees of freedom, i.e. *χ* angles in this case), while having fewer troubles dealing with bulky amino acids with smaller number of degrees of freedom. New training methods that might improve assignment of side chains for these amino acids is an area of further research interest.

**Table 5:**
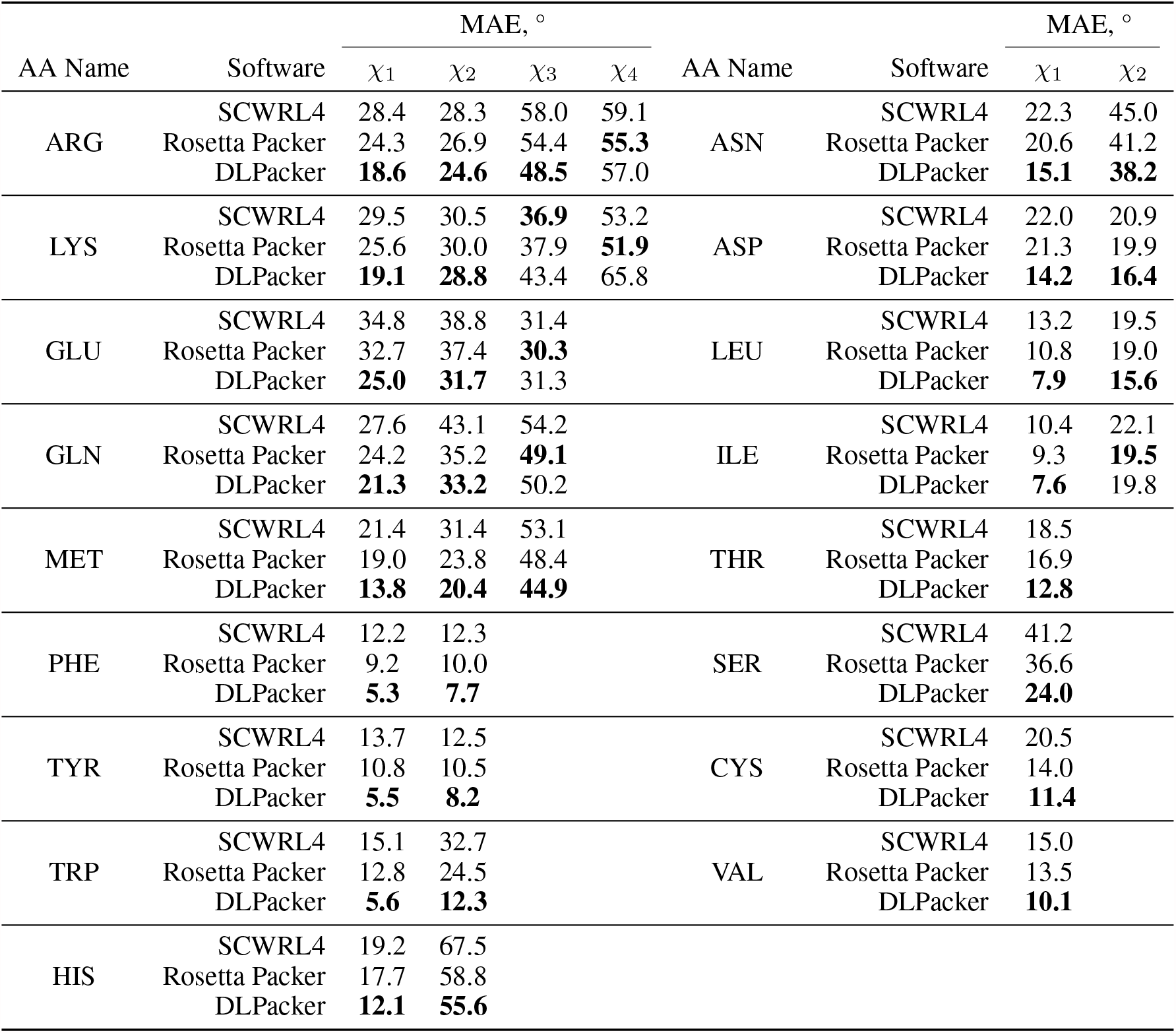
Comparison between Mean Absolute Errors of *χ* angles for SCWRL4, Rosetta Packer and DLPacker. The values were obtained by averaging MAE values for ∼1,000 PDB structures from the validation set.

| AA Name | Software | MAE, $^{\circ}$ | | | | AA Name | Software | MAE, $^{\circ}$ | |
| --- | --- | --- | --- | --- | --- | --- | --- | --- | --- |
| | | $\chi_1$ | $\chi_2$ | $\chi_3$ | $\chi_4$ | | | $\chi_1$ | $\chi_2$ |
| ARG | SCWRL4 | 28.4 | 28.3 | 58.0 | 59.1 | ASN | SCWRL4 | 22.3 | 45.0 |
|  | Rosetta Packer | 24.3 | 26.9 | 54.4 | **55.3** |  | Rosetta Packer | 20.6 | 41.2 |
|  | DLPacker | **18.6** | **24.6** | **48.5** | 57.0 |  | DLPacker | **15.1** | **38.2** |
| LYS | SCWRL4 | 29.5 | 30.5 | **36.9** | 53.2 | ASP | SCWRL4 | 22.0 | 20.9 |
|  | Rosetta Packer | 25.6 | 30.0 | 37.9 | **51.9** |  | Rosetta Packer | 21.3 | 19.9 |
|  | DLPacker | **19.1** | **28.8** | 43.4 | 65.8 |  | DLPacker | **14.2** | **16.4** |
| GLU | SCWRL4 | 34.8 | 38.8 | 31.4 |  | LEU | SCWRL4 | 13.2 | 19.5 |
|  | Rosetta Packer | 32.7 | 37.4 | **30.3** |  |  | Rosetta Packer | 10.8 | 19.0 |
|  | DLPacker | **25.0** | **31.7** | 31.3 |  |  | DLPacker | **7.9** | **15.6** |
| GLN | SCWRL4 | 27.6 | 43.1 | 54.2 |  | ILE | SCWRL4 | 10.4 | 22.1 |
|  | Rosetta Packer | 24.2 | 35.2 | **49.1** |  |  | Rosetta Packer | 9.3 | **19.5** |
|  | DLPacker | **21.3** | **33.2** | 50.2 |  |  | DLPacker | **7.6** | 19.8 |
| MET | SCWRL4 | 21.4 | 31.4 | 53.1 |  | THR | SCWRL4 | 18.5 |  |
|  | Rosetta Packer | 19.0 | 23.8 | 48.4 |  |  | Rosetta Packer | 16.9 |  |
|  | DLPacker | **13.8** | **20.4** | **44.9** |  |  | DLPacker | **12.8** |  |
| PHE | SCWRL4 | 12.2 | 12.3 |  |  | SER | SCWRL4 | 41.2 |  |
|  | Rosetta Packer | 9.2 | 10.0 |  |  |  | Rosetta Packer | 36.6 |  |
|  | DLPacker | **5.3** | **7.7** |  |  |  | DLPacker | **24.0** |  |
| TYR | SCWRL4 | 13.7 | 12.5 |  |  | CYS | SCWRL4 | 20.5 |  |
|  | Rosetta Packer | 10.8 | 10.5 |  |  |  | Rosetta Packer | 14.0 |  |
|  | DLPacker | **5.5** | **8.2** |  |  |  | DLPacker | **11.4** |  |
| TRP | SCWRL4 | 15.1 | 32.7 |  |  | VAL | SCWRL4 | 15.0 |  |
|  | Rosetta Packer | 12.8 | 24.5 |  |  |  | Rosetta Packer | 13.5 |  |
|  | DLPacker | **5.6** | **12.3** |  |  |  | DLPacker | **10.1** |  |
| HIS | SCWRL4 | 19.2 | 67.5 |  |  |  |  |  |  |
|  | Rosetta Packer | 17.7 | 58.8 |  |  |  |  |  |  |
|  | DLPacker | **12.1** | **55.6** |  |  |  |  |  |  |

### 3.3 Effect of Restoration Order

Since our approach implies the restoration of side chains structure in a sequential manner (one amino acid at a time), we need heuristic arguments to choose the most optimal restoration order. In this work, we explored three different ordering strategies. First, we tried a naive strategy that restores side chains in the order of protein’s sequence: from N-terminus to C-terminus. This strategy is both simple and effective since amino acids that are next to each other in the sequence will also be next to each other in the 3D structure. Next, we tried a strategy that orders amino acids according to the number of atoms in their microenvironment and restores side chains in the most crowded microenvironments first. This strategy assumes that it is easier to correctly predict side chain conformation in more crowded microenvironment because there are more interactions and a larger fraction of the volume is already occupied by backbone atoms. It roughly corresponds to first restoring the side chains in the protein’s interior and then gradually moving to its exterior. The last strategy orders amino acids according to the prediction quality and is a two-stage process. First, predictions are made for each amino acid (without actually restoring the side chains) and then residues are sorted by their prediction score, normalized by the number of atoms in a side chain (see **Side Chain Restoration** section). The rationale is that the lower fitness score corresponds to the smaller prediction uncertainty and more likely to lead to correct predictions. Side chains are then restored in the second pass.

All three methods yield similar results, with the first strategy performing slightly worse than the other two for most amino acids (Figure 5). The best performing strategy is the third one (by the prediction score) with the only drawback being that it uses twice as much time and computational resources since we need to pass through the whole protein’s sequence twice. The second strategy (by the number of atoms in the microenvironment) seems to be a perfect compromise between speed and quality.

**Figure 5:**
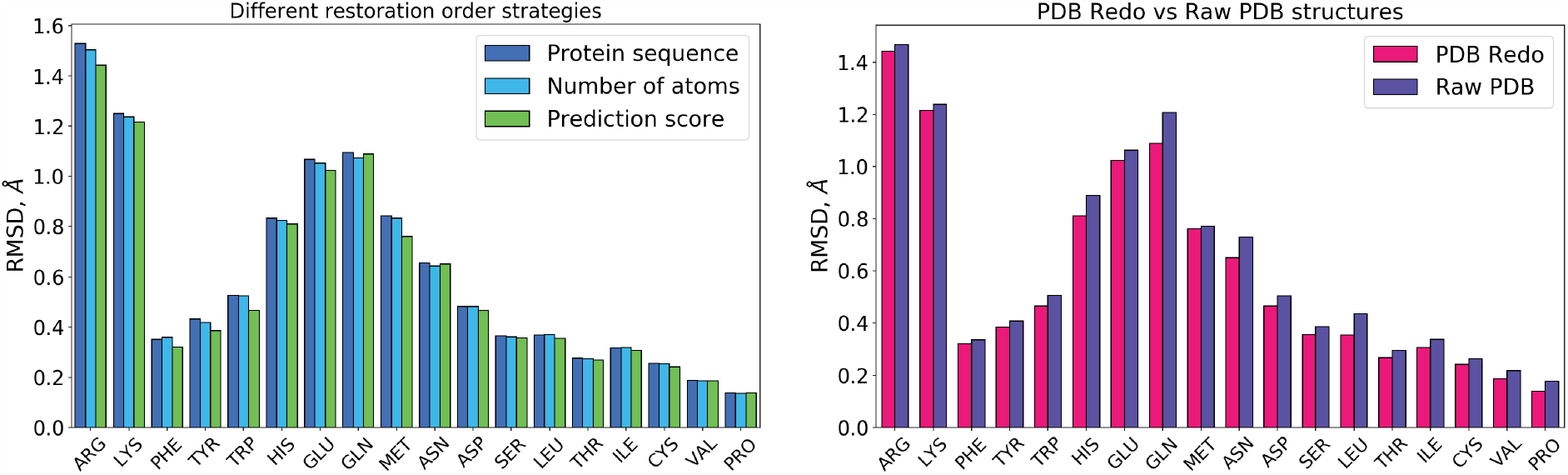
**Left:** Effect of different restoration order on reconstruction RMSD. All three ordering strategies - by sequence, by number of atoms in the microenvironment and by quality of prediction (prediction score) - yield approximately the same results, but latter is the best. **Right:** Comparison between performance on PDB Redo structures and raw PDB structures. Performance on raw PDB structures only slightly worse. More details are in the text.

### 3.4 Performance on Raw PDB Structures

Since our algorithm was trained and validated on PDB structures refined by the PDB Redo server, we wanted to test how well it generalizes to raw PDB structures. [25] The results are shown in Figure 5. The resulting RMSD values are only slightly larger in the case of Raw PDB structures. It is not clear, however, if this difference is due to PDB Redo structures having higher quality (better resolution) or due to the Raw PDB structures just being slightly different from the data our algorithm was trained on.

## 4 Discussion

We developed and comprehensively evaluated a new Deep Learning approach for amino acid side chain prediction in proteins. Besides much better performance compared to other commonly used tools, our approach has a number of other advantages. It can be used on any protein structure, including structures of protein complexes, structures containing DNA, RNA or any small molecules. Our approach can also be utilized in a combination with other physics-based approaches, for example as one component of a scoring function. The output from the neural network is a 3D density map, in which values in every voxel are proportional to probability of finding an amino acid’s side chain atom at that location (Figure 1c), which can be used to score every possible rotamer of an amino acid. This score can then be applied, for example, as a Context Independent One Body Energy term and added to the default Rosetta scoring function. In our experiments, however, this did not lead to further improvement compared to our algorithm (data not shown). There are several other directions that can be explored by connecting our method with existing approaches in order to optimize the whole process of amino acid side chain structure prediction in proteins.

Traditional methods of packing of amino acid side chains rely heavily on hand-crafted scoring functions, which try to take into account all known interactions in proteins. [40] This approach, however, is fundamentally limited by our knowledge of aforementioned interactions. The list of currently known interactions in proteins is quite long. In addition to well known and studied ones, like Van der Waals forces, electrostatic interactions and hydrogen bonds, there is also a list of less studied interactions, which include *π* − *π* stacking, *π*-cation, *π*-anion, *π*-sulphur, NH-*π*, CH-*π*, C-halogen-*π* interactions. [41, 42, 43, 44, 45, 46, 47, 48, 49, 50, 51] To make things more complicated, while all the interactions listed above are usually viewed and characterized as two-body interactions, that is just a simplification of the real picture and many interactions can be highly affected by multi-body effects. Other still unknown two-body and more complicated multi-body interactions are also likely to exist in proteins. [52, 53, 54, 55] All these factors enormously complicate the development of scoring functions for protein packing and make it extremely difficult to estimate the strengths of all interactions and their dependence on geometry and nature of the microenvironment.

Deep Learning approaches, on the other hand, have no need for explicitly defined scoring functions and do not require any assumptions on strength and nature of any existing interactions. [8, 56, 57, 58, 59] Studies suggest that current number of structures in PDB allows for careful and precise estimation of interactions between amino acids, including multi-body interactions. [53, 55] Trying to elucidate the roots of DLPacker’s performance, we looked at some of the structures where it yielded correct prediction, while Rosetta Packer (the better performing of two algorithms we compared to DLPacker to in this work) did not. Results are presented in Figure 6. Visual inspection shows that the most frequent reason of Rosetta Packer’s failure was positioning side chains facing outwards (into solution) instead of burying into a protein. This is true for the most structures shown in Figure 6 - instead of putting the target side chain in position to interact with other side chains shown, Rosetta Packer put it facing outwards and often formed no strong interactions at all. This may be a result of either underestimation of the strengths of some interactions or drawing from a limited number of initial rotamers which do encompass the correct configuration of side chain atoms. In our experiments experiments, however, further increase of the number of initial rotamers only marginally improved RMSD, while drastically increasing computational time.

**Figure 6:**
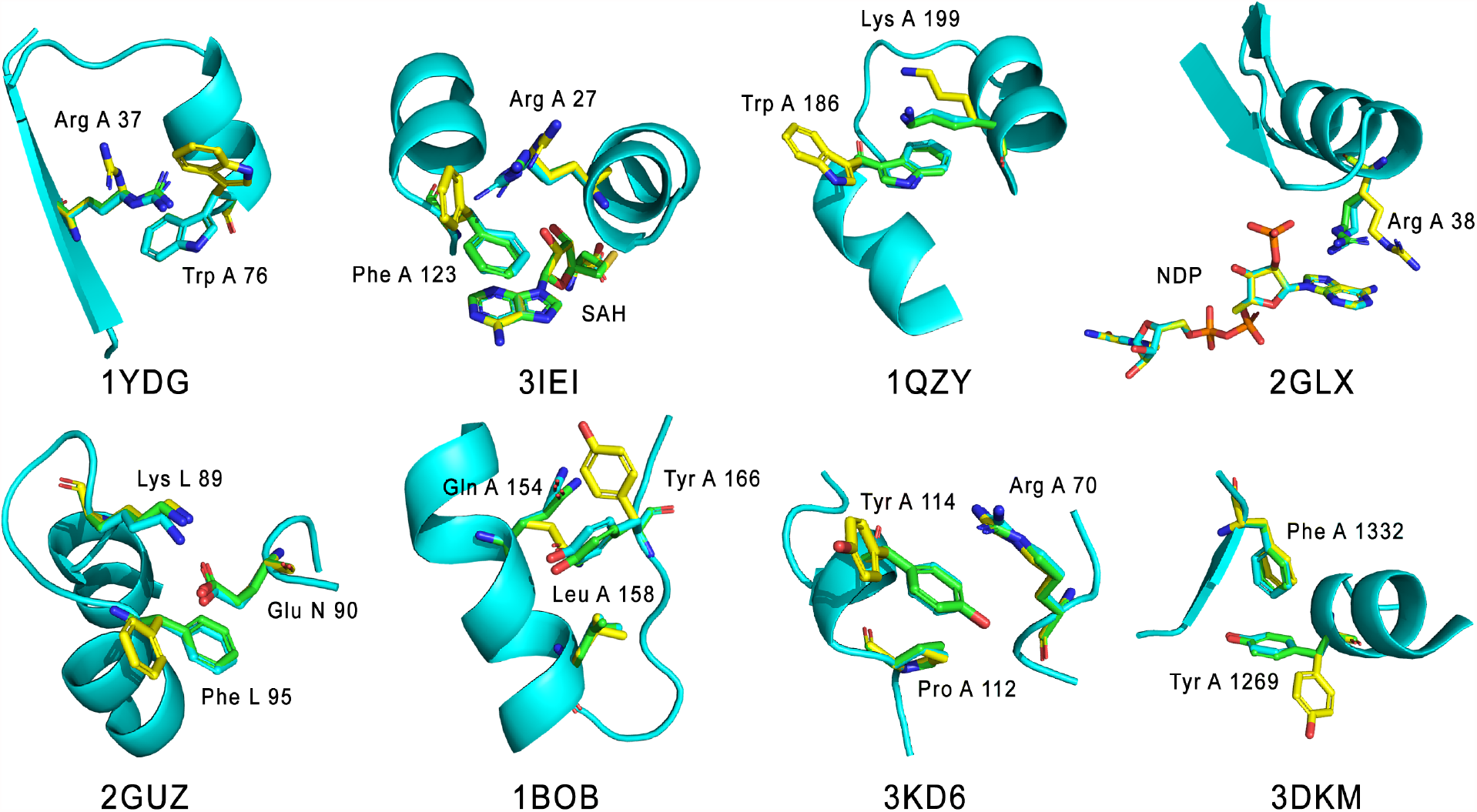
Examples of amino acids for which DLPacker yielded correct result, but Rosetta Packer did not. Backbone is shown in cartoon representation (cyan) and selected side chains are shown using stick representation. Ground truth side chains are shown in green, DLPacker’s output in cyan and Rosetta Packer’s in yellow. PDB codes of structures and amino acid names are shown as well.

We hypothesize that the source of the DLPacker’s high performance is the ability to carefully infer strengths of multiple two-body and multi-body interactions existing in proteins. We note that DLPacker demonstrates the largest improvement for aromatic amino acids (Phe, Tyr and Trp). These amino acids are known to form the largest number of various X-*π* interactions, which involve their aromatic systems. These interactions are also likely to be highly influenced by the amino acid’s microenvironment due to high polarizability of aromatic electrons, potentially leading to complicated dependencies on geometry and nature of its microenvironment.

As was already discussed, our algorithm outperforms other existing methods such as SCWRL4 and Rosetta Packer. It should be noted, however, that currently this comes at the cost of the increased computational time and resources (GPU). Despite the fact that SCWRL4 shows poorer metrics, it is highly optimized for speed and is notably faster than our approach. In our experiments, SCWRL4 was about seven times faster than out algorithm. Rosetta Packer, on the other hand, was about two times slower than the DLPacker (if run with the maximum number of rotamers to achieve the performance we report in this work). If run with default parameters, Rosetta Packer becomes significantly faster at the cost of worse metrics, which are closer to that of SCWRL4. These arguments suggest that our new method can be effectively utilized for situations that require highly precise side chain localization and where the computational times and resources are not a limiting step. It will be important to develop our approach to make it faster and work in conjunction with other available models as part of an integrated workflow for structural biology and protein engineering applications.

## 5 Conclusions

Structural prediction of side chain orientation necessitates effectively filling space while simultaneously avoiding steric clashes. Here, we employed a 3D convolution network to better capture the underlying three-dimensional spatial relationships and show our novel approach towards amino acid side chain packing outperforms classical methods. While DLPacker translates atomic coordinates to predict a spatial density before restoring the coordinates of the improved side chains, we envision further refinements to our method could remove the necessary spatial translation and simply predict the side chain coordinates from only a list of input 3D coordinates.

## 6 Code Availability

The code for DLPacker with some usage examples is available at https://github.com/nekitmm/DLPacker.

## 7 Acknowledgements

MM and ABK acknowledges the support from the Center for Theoretical Biological Physics sponsored by the NSF (PHY-2019745).

## References

[1] Georgii G Krivov, Maxim V Shapovalov, and Roland L Dunbrack Jr. Improved prediction of protein side-chain conformations with scwrl4. Proteins: Structure, Function, and Bioinformatics, 77(4):778–795, 2009.

[2] Thomas Gaillard, Nicolas Panel, and Thomas Simonson. Protein side chain conformation predictions with an mmgbsa energy function. Proteins: Structure, Function, and Bioinformatics, 84(6):803–819, 2016.

[3] Aleksandra E Badaczewska-Dawid, Andrzej Kolinski, and Sebastian Kmiecik. Computational reconstruction of atomistic protein structures from coarse-grained models. Computational and structural biotechnology journal, 18:162–176, 2020.

[4] Patricia Francis-Lyon and Patrice Koehl. Protein side-chain modeling with a protein-dependent optimized rotamer library. Proteins: Structure, Function, and Bioinformatics, 82(9):2000–2017, 2014.

[5] John M Jumper, Nabil F Faruk, Karl F Freed, and Tobin R Sosnick. Accurate calculation of side chain packing and free energy with applications to protein molecular dynamics. PLoS computational biology, 14(12):e1006342, 2018.

[6] Maxim V Shapovalov and Roland L Dunbrack Jr. A smoothed backbone-dependent rotamer library for proteins derived from adaptive kernel density estimates and regressions. Structure, 19(6):844–858, 2011.

[7] Izhar Wallach, Michael Dzamba, and Abraham Heifets. Atomnet: a deep convolutional neural network for bioactivity prediction in structure-based drug discovery. arXiv preprint 1510.02855, 2015.

[8] Andrew W Senior, Richard Evans, John Jumper, James Kirkpatrick, Laurent Sifre, Tim Green, Chongli Qin, Augustin žídek, Alexander WR Nelson, Alex Bridgland, et al. Improved protein structure prediction using potentials from deep learning. Nature, 577(7792):706–710, 2020.

[9] Wen Torng and Russ B Altman. 3d deep convolutional neural networks for amino acid environment similarity analysis. BMC bioinformatics, 18(1):1–23, 2017.

[10] Joseph M Cunningham, Grigoriy Koytiger, Peter K Sorger, and Mohammed AlQuraishi. Biophysical prediction of protein–peptide interactions and signaling networks using machine learning. Nature methods, 17(2):175–183, 2020.

[11] Ethan C Alley, Grigory Khimulya, Surojit Biswas, Mohammed AlQuraishi, and George M Church. Unified rational protein engineering with sequence-based deep representation learning. Nature methods, 16(12):1315–1322, 2019.

[12] Ken Nagata, Arlo Randall, and Pierre Baldi. Sidepro: A novel machine learning approach for the fast and accurate prediction of side-chain conformations. Proteins: Structure, Function, and Bioinformatics, 80(1):142–153, 2012.

[13] Łukasz Maziarka, Tomasz Danel, Sławomir Mucha, Krzysztof Rataj, Jacek Tabor, and Stanislaw Jastrze?bski. Molecule attention transformer. arXiv preprint 2002.08264, 2020.

[14] Yipin Lei, Shuya Li, Ziyi Liu, Fangping Wan, Tingzhong Tian, Shao Li, Dan Zhao, and Jianyang Zeng. Camp: a convolutional attention-based neural network for multifaceted peptide-protein interaction prediction. bioRxiv, 2020.

[15] Alexander Rives, Joshua Meier, Tom Sercu, Siddharth Goyal, Zeming Lin, Jason Liu, Demi Guo, Myle Ott, C Lawrence Zitnick, Jerry Ma, et al. Biological structure and function emerge from scaling unsupervised learning to 250 million protein sequences. Proceedings of the National Academy of Sciences, 118(15), 2021.

[16] John Ingraham, Vikas Kamur Garg, Regina Barzilay, and Tommi S Jaakkola. Generative models for graph-based protein design. 2021.

[17] Surojit Biswas, Grigory Khimulya, Ethan C Alley, Kevin M Esvelt, and George M Church. Low-n protein engineering with data-efficient deep learning. Nature Methods, 18(4):389–396, 2021.

[18] John Jumper, Richard Evans, Alexander Pritzel, Tim Green, Michael Figurnov, Olaf Ronneberger, Kathryn Tunyasuvunakool, Russ Bates, Augustin Žídek, Anna Potapenko, et al. Highly accurate protein structure prediction with alphafold. Nature, pages 1–11, 2021.

[19] Gang Xu, Qinghua Wang, and Jianpeng Ma. Opus-rota3: Improving protein side-chain modeling by deep neural networks and ensemble methods. Journal of Chemical Information and Modeling, 60(12):6691–6697, 2020.

[20] Ke Liu, Xiangyan Sun, Jun Ma, Zhenyu Zhou, Qilin Dong, Shengwen Peng, Junqiu Wu, Suocheng Tan, Günter Blobel, and Jie Fan. Prediction of amino acid side chain conformation using a deep neural network. arXiv preprint 1707.08381, 2017.

[21] Ke Liu, Zekun Ni, Zhenyu Zhou, Suocheng Tan, Xun Zou, Haoming Xing, Xiangyan Sun, Qi Han, Junqiu Wu, and Jie Fan. Molecular modeling with machine-learned universal potential functions. arXiv preprint 2103.04162, 2021.

[22] Chen Yanover, Ora Schueler-Furman, and Yair Weiss. Minimizing and learning energy functions for side-chain prediction. Journal of Computational Biology, 15(7):899–911, 2008.

[23] Faruck Morcos, Andrea Pagnani, Bryan Lunt, Arianna Bertolino, Debora S Marks, Chris Sander, Riccardo Zecchina, José N Onuchic, Terence Hwa, and Martin Weigt. Direct-coupling analysis of residue coevolution captures native contacts across many protein families. Proceedings of the National Academy of Sciences, 108(49):E1293–E1301, 2011.

[24] Raghav Shroff, Austin W Cole, Daniel J Diaz, Barrett R Morrow, Isaac Donnell, Ankur Annapareddy, Jimmy Gollihar, Andrew D Ellington, and Ross Thyer. Discovery of novel gain-of-function mutations guided by structure-based deep learning. ACS Synthetic Biology, 9(11):2927–2935, 2020.

[25] Robbie P Joosten, Jean Salzemann, Vincent Bloch, Heinz Stockinger, A-C Berglund, Christophe Blanchet, Erik Bongcam-Rudloff, Christophe Combet, Ana L Da Costa, Gilbert Deleage, et al. Pdb_redo: automated re-refinement of x-ray structure models in the pdb. Journal of applied crystallography, 42(3):376–384, 2009.

[26] Vladimir Iglovikov and Alexey Shvets. Ternausnet: U-net with vgg11 encoder pre-trained on imagenet for image segmentation. arXiv preprint 1801.05746, 2018.

[27] Zongwei Zhou, Md Mahfuzur Rahman Siddiquee, Nima Tajbakhsh, and Jianming Liang. Unet++: A nested u-net architecture for medical image segmentation. In Deep learning in medical image analysis and multimodal learning for clinical decision support, pages 3–11. Springer, 2018.

[28] Olaf Ronneberger, Philipp Fischer, and Thomas Brox. U-net: Convolutional networks for biomedical image segmentation. In International Conference on Medical image computing and computer-assisted intervention, pages 234–241. Springer, 2015.

[29] Özgün Çiçek, Ahmed Abdulkadir, Soeren S Lienkamp, Thomas Brox, and Olaf Ronneberger. 3d u-net: learning dense volumetric segmentation from sparse annotation. In International conference on medical image computing and computer-assisted intervention, pages 424–432. Springer, 2016.

[30] Md Zahangir Alom, Chris Yakopcic, Mahmudul Hasan, Tarek M Taha, and Vijayan K Asari. Recurrent residual u-net for medical image segmentation. Journal of Medical Imaging, 6(1):014006, 2019.

[31] Rebecca F Alford, Andrew Leaver-Fay, Jeliazko R Jeliazkov, Matthew J O’Meara, Frank P DiMaio, Hahnbeom Park, Maxim V Shapovalov, P Douglas Renfrew, Vikram K Mulligan, Kalli Kappel, et al. The rosetta allatom energy function for macromolecular modeling and design. Journal of chemical theory and computation, 13(6):3031–3048, 2017.

[32] Hahnbeom Park, Philip Bradley, Per Greisen Jr, Yuan Liu, Vikram Khipple Mulligan, David E Kim, David Baker, and Frank DiMaio. Simultaneous optimization of biomolecular energy functions on features from small molecules and macromolecules. Journal of chemical theory and computation, 12(12):6201–6212, 2016.

[33] Andrew Leaver-Fay, Jack Snoeyink, and Brian Kuhlman. On-the-fly rotamer pair energy evaluation in protein design. In International Symposium on Bioinformatics Research and Applications, pages 343–354. Springer, 2008.

[34] Andrew Leaver-Fay, Brian Kuhlman, and Jack Snoeyink. An adaptive dynamic programming algorithm for the side chain placement problem. In Biocomputing 2005, pages 16–27. World Scientific, 2005.

[35] Andrew Leaver-Fay, Brian Kuhlman, and Jack Snoeyink. Rotamer-pair energy calculations using a trie data structure. In International Workshop on Algorithms in Bioinformatics, pages 389–400. Springer, 2005.

[36] Sidhartha Chaudhury, Sergey Lyskov, and Jeffrey J Gray. Pyrosetta: a script-based interface for implementing molecular modeling algorithms using rosetta. Bioinformatics, 26(5):689–691, 2010.

[37] Jian Peng, Raghavendra Hosur, Bonnie Berger, and Jinbo Xu. itreepack: Protein complex side-chain packing by dual decomposition. arXiv preprint 1504.05467, 2015.

[38] Mohammad Moghadasi, Hanieh Mirzaei, Artem Mamonov, Pirooz Vakili, Sandor Vajda, Ioannis Ch Paschalidis, and Dima Kozakov. The impact of side-chain packing on protein docking refinement. Journal of chemical information and modeling, 55(4):872–881, 2015.

[39] Mark A Olson and Michael S Lee. Structure refinement of protein model decoys requires accurate side-chain placement. Proteins: Structure, Function, and Bioinformatics, 81(3):469–478, 2013.

[40] Steven A Combs, Benjamin K Mueller, and Jens Meiler. Holistic approach to partial covalent interactions in protein structure prediction and design with rosetta. Journal of chemical information and modeling, 58(5):1021–1036, 2018.

[41] Niki Zacharias and Dennis A Dougherty. Cation–π interactions in ligand recognition and catalysis. Trends in pharmacological sciences, 23(6):281–287, 2002.

[42] Justin P Gallivan and Dennis A Dougherty. Cation-π interactions in structural biology. Proceedings of the National Academy of Sciences, 96(17):9459–9464, 1999.

[43] Maria Brandl, Manfred S Weiss, Andreas Jabs, Jürgen Sühnel, and Rolf Hilgenfeld. Ch-π-interactions in proteins. Journal of molecular biology, 307(1):357–377, 2001.

[44] SK Burley and GA Petsko. Amino-aromatic interactions in proteins. FEBS letters, 203(2):139–143, 1986.

[45] Sandeep Kumar and Ruth Nussinov. Close-range electrostatic interactions in proteins. ChemBioChem, 3(7):604– 617, 2002.

[46] KG Tina, Rana Bhadra, and Narayanaswamy Srinivasan. Pic: protein interactions calculator. Nucleic acids research, 35(Suppl_2):W473–W476, 2007.

[47] Xavier Lucas, Antonio Bauzá, Antonio Frontera, and David Quinonero. A thorough anion–π interaction study in biomolecules: on the importance of cooperativity effects. Chemical science, 7(2):1038–1050, 2016.

[48] Ishu Saraogi, VG Vijay, Soma Das, K Sekar, and TN Guru Row. C–halogen… π interactions in proteins: a database study. Crystal engineering, 6(2):69–77, 2003.

[49] Pascal Auffinger, Franklin A Hays, Eric Westhof, and P Shing Ho. Halogen bonds in biological molecules. Proceedings of the National Academy of Sciences, 101(48):16789–16794, 2004.

[50] Rubicelia Vargas, Jorge Garza, David A Dixon, and Benjamin P Hay. How strong is the cα-h…o=c hydrogen bond? Journal of the American Chemical Society, 122(19):4750–4755, 2000.

[51] Vivek Philip, Jason Harris, Rachel Adams, Don Nguyen, Jeremy Spiers, Jerome Baudry, Elizabeth E Howell, and Robert J Hinde. A survey of aspartate-phenylalanine and glutamate-phenylalanine interactions in the protein data bank: Searching for anion-π pairs. Biochemistry, 50(14):2939–2950, 2011.

[52] Silvana Pinheiro, Ignacio Soteras, Josep Lluís Gelpí, François Dehez, Christophe Chipot, F Javier Luque, and Carles Curutchet. Cation–π–cation interactions in structural biology. In BSC Doctoral Symposium (2nd: 2015: Barcelona), pages 103–105. Barcelona Supercomputing Center, 2015.

[53] Kristoffer Enøe Johansson and Thomas Hamelryck. A simple probabilistic model of multibody interactions in proteins. Proteins: Structure, Function, and Bioinformatics, 81(8):1340–1350, 2013.

[54] Iain H Moal, Rocco Moretti, David Baker, and Juan Fernández-Recio. Scoring functions for protein–protein interactions. Current opinion in structural biology, 23(6):862–867, 2013.

[55] Xiang Li and Jie Liang. Geometric cooperativity and anticooperativity of three-body interactions in native proteins. Proteins: Structure, Function, and Bioinformatics, 60(1):46–65, 2005.

[56] Janaina Cruz Pereira, Ernesto Raul Caffarena, and Cicero Nogueira Dos Santos. Boosting docking-based virtual screening with deep learning. Journal of chemical information and modeling, 56(12):2495–2506, 2016.

[57] Chao Shen, Junjie Ding, Zhe Wang, Dongsheng Cao, Xiaoqin Ding, and Tingjun Hou. From machine learning to deep learning: Advances in scoring functions for protein–ligand docking. Wiley Interdisciplinary Reviews: Computational Molecular Science, 10(1):e1429, 2020.

[58] Robin Pearce and Yang Zhang. Deep learning techniques have significantly impacted protein structure prediction and protein design. Current Opinion in Structural Biology, 68:194–207, 2021.

[59] Isabella A Guedes André, MS Barreto, Diogo Marinho, Eduardo Krempser, Mélaine A Kuenemann, Olivier Sperandio, Laurent E Dardenne, and Maria A Miteva. New machine learning and physics-based scoring functions for drug discovery. Scientific reports, 11(1):1–19, 2021.

